# Efficient coding of subjective value

**DOI:** 10.1101/358317

**Authors:** Rafael Polanía, Michael Woodford, Christian C. Ruff

**Affiliations:** Zurich Center for Neuroeconomics (ZNE), Department of Economics, University of Zurich, Switzerland; Decision Neuroscience Lab, Department of Health Sciences and Technology, ETH Zurich, Switzerland; Department of Economics, Columbia University, New York, USA

## Abstract

Preference-based decisions are essential for survival, for instance when deciding what we should (not) eat. Despite their importance, choices based on preferences are surprisingly variable and can appear irrational in ways that have defied mechanistic explanations. Here we propose that subjective valuation results from an inference process that accounts for the information structure of values in the environment and that maximizes information in value representations in line with demands imposed by limited coding resources. A model of this inference process explains the variability in both subjective value reports and preference-based choices, and predicts a new preference illusion that we validate with empirical data. Interestingly, the same model also explains the level of confidence associated with these reports. Our results imply that preference-based decisions reflect information-maximizing transmission and statistically optimal decoding of subjective values by a limited-capacity system. These findings provide a unified account of how humans perceive and valuate the environment to optimally guide behavior.

## Introduction

At any given moment, organisms receive much more sensory and interoceptive information than they can physically process. These capacity limitations are thought to have biased brain evolution towards information-processing strategies that are maximally efficient for the control of behavior, an idea known as *efficient coding*^1,2^. Such efficient-coding strategies can be observed in sensory systems where the precision with which neural representations encode different states of an environmental variable (e.g., different orientations) is proportional to the frequency with which this state is actually encountered^3,4^. This strategy ensures that the information encoded is as great as possible given the dynamic range of the physical system used to represent it^5^. However, these representations not only need to be efficiently encoded but also need to be decoded and interpreted so that the resulting percepts provide maximally accurate information about the true state of the world and the organism. Bayesian statistics imply that optimal perceptual processes would have to combine the representation of the environmental information (i.e., the likelihood of a state) with an a-priori expectation of these states^6–8^. While efficient coding and Bayesian decoding theories may appear related, they have only recently been combined in a unified theoretical framework that can account for various low-level perceptual biases^4,9^. But whether similar encoding and decoding strategies also operate in other domains than low-level perceptual systems remains an open question.

In the domain of preference-based decisions, it is commonly assumed that organisms rely on strategies that maximize the utility of the chosen option, based on stable and accurate representations of preferences that are not systematically affected by processing-resource constraints. However, empirically-observed choice behavior often deviates from the predictions of rational choice theory^10^. Purely descriptive theories of such anomalies have been offered, postulating either competition between parallel action-selection processes based on simple heuristics^10^ or some type of arbitrary external noise that has no clear psychological or neural basis^11–13^. While such theories can account for some observed effects of choice variability, biases, or confidence in isolation^11,13–16^, a common framework linking these different aspects of behavior is largely missing. Moreover, these models sometimes contain assumptions about value representations that appear implausible given the constraints imposed by the limited-capacity nature of biological systems.

In order to account for these limitations, recent work has sought to find shared principles in the mechanisms underlying subjective valuation and sensory perception^11,13,16–22^. Theories from this line of research have suggested that subjective value representations may resemble percepts in that they are derived by inference processes that exploit prior information about the relevant distribution of value stimuli in the environment^17,18,20,22,23^. Moreover, related lines of work suggest that neural reward circuits can flexibly adapt to different value contexts in the evironment^24–27^, possibly consistent with the notion that neural resources are allocated efficiently to the encoding of subjective values. However, it is unknown whether efficient coding and Bayesian decoding principles are indeed used jointly to generate preference-representations, and whether this information-processing scheme can explain the variability, biases, and confidence in value-based decisions in humans. This lack of knowledge may reflect that the distribution of subjective values in the natural environment is not easily measurable (in contrast to corresponding distributions of sensory signals^28^), since it depends on the long-term experience of each specific organism with the objects in its environment^27^.

Here we propose a way to test directly whether preference-based decisions are indeed guided by a value representation scheme that combines both efficient coding and Bayesian-decoding principles. We achieve this by introducing a novel approach for studying subjective valuation that takes account of the important fact that neither decision makers nor experimenters have direct access to “true” subjective values underlying all value-related behaviors (e.g., value ratings, value-based choices, etc.). We demonstrate with modelling and behavioral experiments that choice variability, biases, and confidence in human preference-based decisions can all be explained by a single value-inference process. This process maximizes information transmission by optimally allocating limited resources to value representations, based on prior knowledge about the natural distribution of object values in the individual environment. Our approach accounts comprehensively for several aspects of value judgements and value-based choices, proposes that humans may make value-based choices optimally given resource constraints, and provides a unified perspective on how the brain may employ common principles to guide behavior on the basis of both perceptual and value information.

## Results

### Efficient coding of subjective value

In studies of perceptual decisions, experimenters usually have complete knowledge of any experimental stimulus value *v* (for instance, the angular orientation of displayed Gabor patches). This is different in experiments studying value-based choices, since experimenters have no direct access to the “true” *v* value of the presented object to an observer (Fig. 1a). Here we assume that this “true” value *v* has been shaped by each observer’s personal history of experiences with this type of object and is therefore entirely subjective. A common strategy adopted by experimenters is thus to first derive an estimate 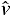 of this subjective value – based on empirical choices or subjective reports (Fig. 1a) – that is subsequently used as input to a decision model^11,13–15,29–31^. However, we will show that this strategy is suboptimal because the value estimates 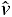 are likely to be as inaccurate and biased as the subsequent choices. This is because the observer herself does not have direct access to the “true” value *v* – after all, she does not have perfect memory of all her lifetime experiences. Thus, the observer needs to derive an *estimate* of the object’s value *v* every time this is necessary, for instance when having to rate this value or when choosing between this object and another one. Any noise and bias resulting from encoding/decoding processes used to infer this value estimate should thus affect any type of behavior in similar ways. Given these limitations, we elaborate a new approach that yields more precise estimates of subjective values for the study of preference-based decisions, based on the principles of efficient coding and Bayesian decoding.

**Figure 1.**
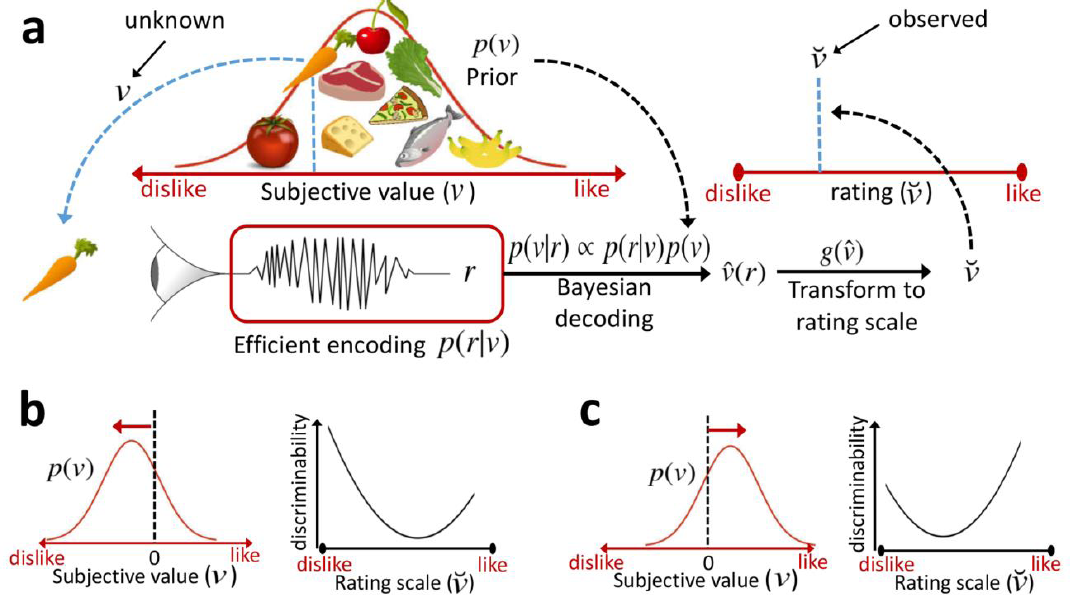
Simplified scheme of the value inference model. **a)** In this example, the prior belief *p(v)* matches the distribution of subjective values *v* of supermarket products. When asked to rate an item, the likelihood function *p(r∣v)* is constrained by the prior via efficient coding, where *r* is the internal response given the subjective value *v*. The prior is combined with the likelihood to generate a posterior distribution *p(r∣v)* via Bayes rule, to generate a subjective value estimate 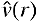. This estimate is subsequently mapped to the bounded rating scale imposed by the experimenter, resulting in an observed rating 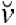. Crucially, unlike for experiments of perception, the experimenter has no access to the “true” stimulus value *v* that the participant uses to generate a rating 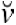. For easier readability, in this scheme we have omitted external sources of noise. **b,c)** Choice consistency predictions as a function of the prior and the rating scale position. Prior distributions with higher density over low subjective values lead to higher choice accuracy for low-valued goods (panel b); on the other hand, prior distributions with higher density over high subjective values lead to higher choice accuracy for higher valued goods (panel c).

We model valuation as a probabilistic inference process incorporating both encoding and decoding (Fig. 1a; see methods for full details). Presentation of an object with “true” stimulus value *v* elicits an internal noisy response *r* (*encoding*) that is used by the observer to generate a subjective value estimate 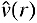 (*decoding)* that is reported behaviorally (Fig. 1a). In experimental settings, such behavioral reports typically have to be given on physically bounded rating scales^11,13–15,29^ that can differ across different settings. To account for this step, we assume that the individual’s internal subjective scale for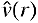 is physically unbounded but can be flexibly mapped to any bounded scale in line with experimental demands. We assume that this is achieved by a function 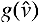 that preserves monotonicity, thus allowing a one-to-one mapping from 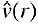 on the internal subjective scale to the response 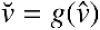on the specific physical response scale employed in an experiment.

Inspired by previous work in the perceptual domain^9^, we assume that *encoding* of subjective values is efficient in the sense that the mutual information between the stimulus values *v* and the internal response *r* is maximized. This results in optimal use of the underlying neuronal scale given the expected/learned natural distribution of values in the given environment (i.e., the prior). We assume that the (noisy) internal response is given by 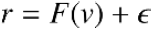, with an error term 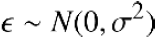 that quantifies the degree of noise in any internal value representation (i.e., the term σ is constant for all possible values of *F(v)*). Different from work in the perceptual domain and standard approaches in neuroeconomic studies, we suppose that the experimenter has no knowledge about the specific stimulus value *v*_0_ that generated the reported rating scale value 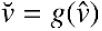. However, if the experimenter obtains several value ratings 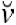for a given good (and if she repeats this for the full distribution of goods in a given context or environment), then it is possible to infer the stimulus value *v*_0_ that is most likely to have generated the observed rating distribution *p(v)* for that good. Parameterizing the prior distribution by a set of parameters, the distribution of estimates (observed ratings in the experiment) for a given good can thus be derived as follows (see methods for details):

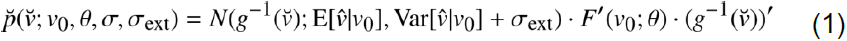

Where 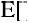 and 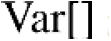 are the expected value and variance of the subjective value 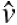 given the input value *v0*, and σ_ext_ is some unspecific noise in decision process (see methods and Supplementary Note 1). Under certain assumptions about the parametric forms of the prior *p(v;θ)* and *g(v)*, an analytical expression for Eq. 1 can be derived (see methods). Thus, for any experimental data set comprising *N* value ratings 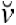 for each of *M* goods sampled randomly from all goods encountered in a given environment (and that therefore follow *p(v;θ)*), one can find the prior *θ*, the internal valuation noise σ, and the “true” values *v(1,…,M)* that maximize the likelihood of the observed set of ratings 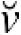, under the constraint that *v(1,…,M)* is distributed according to *p(v;θ)*(Supplementary Fig. 1). Thus, this efficient coding model can generate the observed trial-to-trial variability in observed value ratings by sampling from the distribution 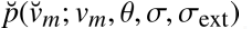.

If individuals employ the efficient-coding approach described above, then the values *v(1,…,M)* obtained by maximizing the likelihood 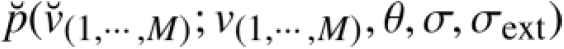using the rating data should predict subsequent choice behavior in a multi-choice task between the same goods *M*. For instance, in a two-alternative choice task, the optimal strategy is to choose good 1 over good 2 if 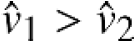, where the value estimates 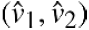are the Bayesian posterior means of each good, respectively (see methods).

One can also make predictions of choice discriminability between two objects based on their position in the rating scale (Fig. 1b,c). If the prior distribution has higher density over low subjective values, then predicted discriminability resembles a u-shaped function, but with higher choice accuracy for lower valued goods (Fig. 1b). On the other hand, if the prior distribution has higher density over high subjective values, then higher choice accuracy should be observed for higher valued goods (Fig. 1c). Interestingly, the latter prediction would be diametrically opposite to predictions based on Weber’s law^8^,which generally assumes that higher value magnitudes should lead to poorer discrimination. Our efficient coding theory implies that Weber’s Law should hold in the case of a particular kind of prior distribution that may be realistic for some sensory magnitudes, but not for the kind of distribution of values for consumption goods assumed here.

### Subjective value rating and choice behavior

In a first behavioral experiment (Experiment 1), we presented a set of food items to n=38 study participants and asked them to indicate on a continuous rating scale their preference to consume the presented item (Fig 2a; methods). Crucially, the participants were familiar with the food products and had seen all of them before the ratings took place, ensuring that they could effectively use the full range of the rating scale. The products (M=64 goods) were a representative sample of products typically encountered in the two biggest chains of supermarkets in Switzerland (participants were aware of this information; see methods). Nevertheless, we ensured prior to testing that participants were indeed familiar with all products.

**Figure 2.**
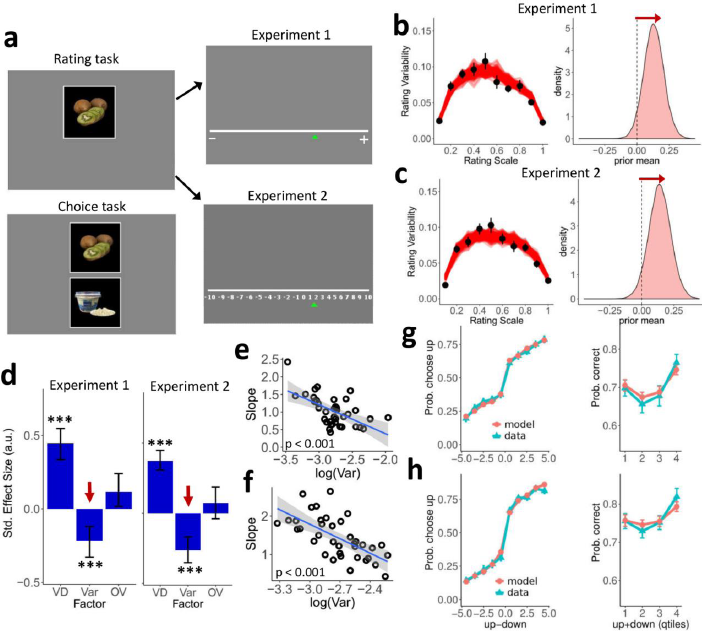
Paradigm and results for Experiments 1 and 2. **a)** Example display from the preference rating phase (two rounds) during which the participants rated their preference to eat the displayed food item using a continuous scale (Experiment 1) or a discrete scale with numerical cues (Experiment 2). The lower-left panel shows an example display from the decision-making task requiring participants to choose which of the two items (upper or lower) they preferred to consume after the experiment. **(b)** Left panels show rating variability plotted as a function of each item’s mean rating across both rounds for experiment 1 (black dots; error bars represent the s.e.m. across participants). We fitted the likelihood function (Eq. 1) to the empirical ratings. To test the quality of the model fits, we simulated 500 experiments in which we draw N=2 ratings for each good and plot the simulated rating variability as a function of the mean rating (semi-transparent red lines, derived exactly as for the empirical data). The rating variability of the simulated experiments is highly consisted with the empirical data. Right panels show that posterior estimates of the expected value of the prior are shifted towards higher rating values (the zero position maps to the center of the rating scale). Based on this shift, our model predicts a discriminability pattern similar to figure 1c. **c)** Same as panel b but for experiment 2. **d)** Standardized estimates from multiple logistic regression show that the higher the value difference VD between the mean ratings, the more consistent the choices. Crucially we also found that the higher the variability 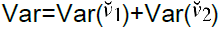 in the ratings of the alternatives, the less consistent the decisions. Total value OV=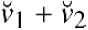 had no reliable influence on choices across the two experiments. This suggests that preference-based choices depend systematically on the properties of internal noisy coding of value. Error bars in this panel represent the 95% highest density interval of the posterior estimates, (***) P<0.001. **e,f)** The trial-to-trial effect shown in panel d was also reflected across participants, as the general level of variability in the rating task correlated negatively with the slope of a logistic regression across participants (experiment 1: panel **e**, βrobust = −0.77±0.20,P<0.001; experiment 2: panel **f**, β_robust_ = −1.2±0.25, P<0.001). **g,h)** Plots of the observed data and the model predictions show that the efficient coding model systematically captures how choice behavior is affected by the two items’ absolute value difference (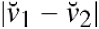, left panels) and overall value (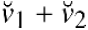 in quartiles, right panels) (panel **g**: experiment 1; panel **h**: experiment 2). The error bars in this panel represent the s.e.m. across participants.

We then asked participants to rate the same items a second time, but crucially, they had been unaware that this second rating phase was going to take place. This was important as it prevented participants from actively memorizing the rating in the first phase, thus allowing a clean estimation of variability in the decoded values (Fig. 2b). We tested whether this variability actually reflected the value coding/decoding operations rather than just random noise or an artifact of the bounded rating scale. To this end, the same participants underwent a series of incentive-compatible choices (see methods) in which they selected from pairs of the previously-rated food items the one item they preferred to eat. We defined a *consistent choice* as a trial in which the subject chose the item they had assigned a higher average rating across the two previous ratings. In line with our assumptions, choice consistency was affected by the value difference between the two items’ prior ratings: The higher the value difference, the more consistent the choices (multiple logistic regression, β_RandEffects_=0.44±0.05, P<0.001; Fig. 2d). This concurs with the long-held notion that stronger evidence leads to more consistent choices^11,13^. Importantly, choice consistency also depended on the variability in the value ratings: The higher this variability for the items on a given trial, the less consistent the decision (βRandEffects = −0.21±0.05, P<0.001; Fig. 2d). Extending this trial-to-trial effect of rating variability, we observed that each participant’s average level of variability in the rating task was negatively correlated with the slope of the logistic regression of her individual choices on the items’ mean value difference (βrobust = −0.77±0.20, P<0.001; Fig.2e). In other words, the higher the variability in the initial value ratings, the less consistent the subsequent choices between the rated items, both compared across trials and across participants. This already suggests that properties of the value coding/decoding operations can somehow affect preference-based choices, but it does not characterize what these properties are and by what mechanisms they may influence the observed decisions. In the next sections, we will address this question by formal tests of the theoretical framework outlined in our model specification.

In Experiment 1, the rating scale was continuous and without numerical cues (Fig. 2a). One may wonder whether rating variability might represent imprecisions in the participants’ assignment of the decoded subjective values to this rating line. We therefore conducted a second experiment (Experiment 2, n=37) in which the rating scale was divided into discrete steps with explicitly assigned numerical values (Fig. 2a; methods). The variability in ratings across this scale clearly resembled the shape observed in Experiment 1 (Fig. 2c) and had a similar significant impact on choice behavior (β_RandEffects_= −0.25±0.05, P<0.001; Fig. 2d). Again, each participant’s level of variability in the rating task correlated negatively with the slope of the regression of choice consistency on value difference between the goods (β_robust_ = −1.2±0.25, P<0.001; Fig. 2f). Thus, the influence of rating variability on subsequent choices does not depend on specifics of the rating procedure but may reflect characteristics of the noisy coding/decoding operations used by the observer to estimate the subjective value. In the next section, we proceed to formally test this notion.

### Testing the efficient coding hypothesis

We now investigate to what extent the observed rating variability in Experiments 1 and 2 can be explained by the efficient coding model (outlined above in methods). We started by finding the set of parameters *v(1,…,M),θ,σ*, and σ_ext_ that maximized the likelihood of the observed set of ratings for each participant, under the constraint that *v(1,…,M*) is distributed following *p(v;θ)* (see Eq. 1 above and methods). In Experiments 1 and 2, the rating data set consisted of *M*=64 and M=61 goods respectively, with *N*=2 ratings for each good. While it may be beneficial to obtain as many ratings as possible for each good (to maximize accuracy in the estimation of subjective values), we limited the number of ratings to *N*=2 in order to prevent participants from actively memorizing previous ratings (methods). The fitted model successfully captured the empirically-observed rating variability (Fig 2b-c) and the distribution of subjective value estimates (Supplementary Fig. 2). We compared the quality of these efficient-coding model fits with those of a simple flexible model that assumes constant Gaussian noise over the rating scale with no prior distribution constraints on the values *v(1,…,M*). For both Experiments 1 and 2, the efficient coding model explained the rating distribution better than the alternative model (Supplementary Fig. 3).

**Figure 3.**
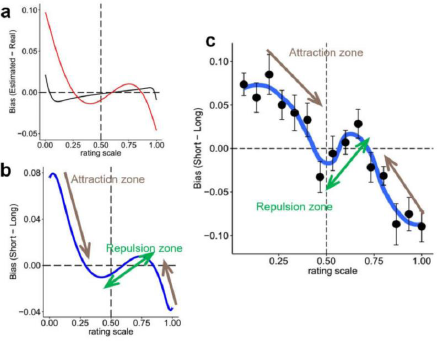
Illusion of preference (Experiment 3). **a)** Model-predicted biases for two degrees of internal measurement noise (σ) during subjective-value estimations: high (red, generated by short valuation exposure) and low (black, generated by long valuation exposure). **b)** Model-predicted differences of the biases for high and low σ across the value rating scale. If the efficient coding hypothesis is correct, both attraction and repulsion should emerge during subjective valuation. **c)** Difference of the empirical estimates between low and high exposure times (data are black dots with s.e.m. across participants, blue line interpolates these data for visualization). Note the qualitative and quantitative overlap with the model prediction in panel b. This suggests that human valuation exhibits complex illusions of subjective preference, as predicted by the Bayesian and efficient coding hypothesis.

Exploration of the rating data revealed that the distribution of ratings was highly variable across participants (Supplementary Figs. 4,5), perhaps indicating that each individual holds different priors over values due to different long-term experience. Moreover, the inferred prior distributions for both experiments revealed that the expected value of the prior across the population was shifted towards higher values (Fig. 2b,c). Choice discriminability should therefore resemble the shape predicted in Figure 1c. If the subjective values of the products are derived using efficient-coding principles, then using the framework described above should allow us to predict each individual’s preference-based decisions. It is important to emphasize that for these prediction analyses, we fixed for each participant the parameters of the prior distribution *θ* and the stimulus values to the specific values obtained when fitting the model to the separate rating data. Using these out-of-sample parameters and values, and only adjusting the encoder and external noise, our model did a good job at predicting the choice data (Fig. 2g,h).

To determine which aspects of the operations formalized in the efficient-coding model are most relevant for explaining behavior, we compared the predictive power of our model (*Model 1*) to that of alternative models (Supplementary Figs. 6,7). In *Model 1.2*, choice predictions were carried out by setting σ = 0 (and only adjusting σ_ext_) in order to investigate whether noise in the subjective value estimates 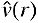due to efficient coding (see Eq. 11 in methods) is relevant for explaining choice behavior. *Model 2* is the standard logit model commonly used in the literature, which simply assumes constant Gaussian noise over the rating scale used for the observed responses 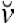. Note that this model is analogous to a Bayesian model with a uniform prior over the rating line. *Model 3* is a variant of the standard logit model that also assumes constant Gaussian noise, but now over the unbounded scale used for the subjective value estimates 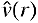. This model is analogous to a Bayesian model with a uniform prior over the unbounded scale. *Model 4* is a Bayesian model that does not assume efficient coding. This model employs Bayesian inference to discover both the “true” stimulus values and the prior distribution, but instead of efficient coding assumes a constant likelihood function over the unbounded scale. For both experiments, the efficient coding model predicted the data best, as evident from visual inspection of the plotted data against the predictions of all models (Supplementary Fig. 6) and from leave-one-out cross-validation metrics (Supplementary Fig. 7).

It may be argued that the specific pattern of observed choices in our experiments could also be captured by a model that does not contain efficient coding but instead fits the full shape of the likelihood function to the observed data, as done e.g. in in previous work on perceptual discrimination^28^. However, it is important to emphasize that our approach does not require complex fits of arbitrary shapes likelihood functions, as these shapes naturally arise from the efficient-coding formalized in our model that only requires fit of one free parameter: Noise in the efficient coding space (for a detailed discussion on this topic see Supplementary Note 2). Thus, the explanatory power of our model does not reflect the general flexibility of Bayesian inference per se but specifically relates to the efficient coding of values embedded in the Bayesian inference process, in close analogy to previous work on low-level sensory perception^9^.

The results presented so far suggest that subjective value representations guiding human preference-based decisions are inferred and employed optimally using both efficient coding and Bayesian decoding. While this is evident in the choices observed in our task, it remains unclear whether internal noise due to efficient coding is the main factor explaining fluctuations and potential biases in subjective value estimations. We investigate this issue in the following section.

### Illusions of subjective value

The theory used here predicts in general that for a stimulus with value *v_0_* near the peak of the prior, the subjective value estimate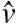 (and the resulting rating 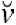) should be biased away from the prior, with the strength of this bias determined by the degree of noise in the internal representations used for inference^9^. Importantly, this repulsion bias runs counter to the predictions of classical Bayesian estimation but has nevertheless been confirmed for basic perceptual tasks (such as orientation and spatial frequency estimation)^4,9^. We thus took a similar approach and investigated whether a conceptually similar type of bias emerges during subjective value estimation, reflecting an expectation-induced preference illusion and further supporting efficient coding of subjective value.

The estimation of this valuation bias necessarily requires knowledge of the exact stimulus value *v_0_* that serves as input on any given trial, which is difficult in our case since the experimenter does not have direct access to *v_0_* (Fig. 1a). In order to cope with this problem, we first derived predictions of the estimation bias for different levels of internal noise σ in the value representations. We assumed that this noise varies with the stimulus presentation times, based on theoretical frameworks postulating that value estimates are gradually constructed using samples from memories/emotions associated with the physical features of the objects^32^. This suggests that a reduction in visual stimulation time should reduce the number of samples that can be drawn and should therefore increase the noise in the internal value representations^33^. To make this intuition explicit, we formulated a mathematical proof confirming that the number of discrete samples (e.g. memories) that can be drawn over time is inversely proportional to the level of encoding noise in a capacity-limited system (Supplementary Note 3). Crucially, this proof provides a normative foundation for the theoretical frameworks^32,34^ motivating our approach and confirms the validity of the assumptions underlying our simulations and experimental strategy.

In order to derive initial qualitative predictions, the prior *p(v)* used for our simulations was obtained based on the results obtained in Experiments 1 and 2, with the consequence that its peak was shifted to the right of the rating scale (Fig 2b,c). The simulations predicted biases for long exposure times (low σ, black line in Figure 3a) and short exposure times (high σ, red line in Fig. 3a) that are markedly different once the value estimates have been mapped on the bounded rating scale via 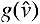. Crucially, the difference between these two predictions (high and low σ) is independent of *v_0_*:

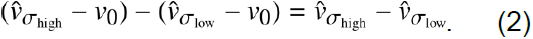

Figure 3b shows that simulated rating trials with short exposure time (i.e., high σ) have stronger repulsive (“anti-Bayesian”) biases near the center of the prior (intersection point to the right of the center of the rating scale), but also stronger attraction biases when *v_0_* is further away from the peak of the prior. For values close to the prior, this prediction agrees with previous work showing repulsive noise-related biases in perceptual tasks^9^; however, as moves away from the prior, our simulations predict the opposite tendency (attraction) that would also be expected based on classical Bayesian frameworks (Supplementary Fig. 8).

We devised a behavioral paradigm (Experiment 3) to investigate whether these model-predicted biases are in fact observed for subjective value estimations. Healthy individuals (n=24) rated goods with similar procedures as in Experiment 1. However, for the first round of ratings, a randomly determined half of the goods for each participant were presented for a duration of 900 ms and the other half for a duration of 2600 ms. In the second round of ratings, the presentation durations were inverted for each good (methods). We then computed the difference in the ratings between short and long durations and plotted this difference as a function of the rating of the long duration (Fig. 3c). The results of this analysis fully match the non-intuitive quantitative predictions of the efficient coding model (Fig. 3b), in showing systematic repulsion for the four data points near the peak of the prior and attraction for the other values that are further away from the prior. Control analyses confirmed that these results are not caused by systematic temporal order effects (no difference between the first and second round of ratings for each of the exposure times; paired t-tests, all P>0.18). Moreover, we also compared the accuracy of our model’s predictions to that of control models in which we factorially varied all possible sources of noise that could in principle have affected the ratings (pre-encoding noise, efficient-coding noise, post-encoding noise, and lapses; see Supplementary Note 1, Supplementary Figure 9). Visual inspection already suggests that the predictions of all control models without efficient coding lead to very different predictions that are not supported by our empirical data (Supplementary Fig. 10). In order to test this quantitatively, we performed a factorial model comparison based on the log factor likelihood ratio approach (LFLR^35^) that quantifies the strength of empirical support given by our data for the presence of each noise source (Supplementary Fig. 9 and Supplementary Table 1). This revealed that the only noise source reliably accounting for the variation in subjective value estimation due to time pressure is internal noise in efficient coding (Bayes Factor > 100; see Supplementary Fig. 9). These findings are not biased by outliers: Inspection of the results for each participant reveals that 21 out of 24 participants in Experiment 3 have a positive LFLR for the internal-noise efficient-coding factor. This strongly suggests that the biases in subjective value estimates observed in Experiment 3 originate in the efficient-coding operations formalized in our model and do not relate to other noise sources commonly postulated in the literature.

### Confidence in subjective valuation

It has been suggested that the perceived confidence in subjective value reports reflects a second-order judgement (the confidence rating) about a first-order judgment (the subjective value rating)^15^. However, two important issues have remained unaddressed. First, previous work has not explicitly defined a generative model of the encoding and decoding of value representations; second-order statements about these ratings would therefore be subject to the same problems curtailing the validity of the ratings themselves (see “Efficient coding of subjective value” section above). Second, previous work has remained agnostic about both the information structure of values in the environment and capacity limitations. We therefore tested whether the reported confidence in subjective value estimations can be predicted based on the encoding/decoding process proposed here. We examined this for two definitions of confidence. The first derives confidence as the second-order judgement of the decoded posterior distribution on the rating scale, as assumed in previous work^15^ (for details see description in Supplementary Table 2). The second assumes that confidence reflects the probability that the rating is “correct”^36,37^(note that a rating task does not have an objectively correct answer, but we can quantify the probability that the rating is correct for a specific observer given her prior and internal noise^38^).

We conducted a new experiment (Experiment 4) in which participants provided value ratings as in experiment 1, but now also gave a confidence rating after each value rating (Fig. 4a). We again derived the subjective values *v(1,…,M*) by maximizing the likelihood function of equation 1 based on the rating data, exactly as for Experiments 1 and 2. We replicated for this new dataset three key results also obtained in experiments 1 and 2: First, the shape of the rating variability across different value levels; second, the accuracy of the model in predicting the rating levels and their variability; third, the shift of the expected prior (i.e., its mean) towards higher values (Fig. 4a-c). We also observed that rating variability was higher for low-rated items (hierarchical linear regression rating variability for each item vs. mean rated value: β_RandEffects_= −0.18±0.04, P<0.001, see Figure4b). Based on all this information, we derived three qualitative predictions for the confidence ratings based on the definitions of confidence formulated above. First, confidence should be higher for rating values near the extremes of the rating scale (Fig. 4d), reflecting the transformation from the subjective space to the bounded scale. This prediction is in line with previous work^15^. Second, given the shift of the prior density towards higher subjective values (Fig. 4c), the efficient coding framework predicts that the second-order judgement of the posterior probability perceived in the rating scale should decrease for item values towards the right side of the rating scale; confidence reports should therefore be higher for items with higher subjective value (Fig. 4d). Third, because lower levels of variability in the rating estimates generate narrower posterior distributions, the average variability of each participant’s ratings should be negatively related to her general level of confidence.

**Figure 4.**
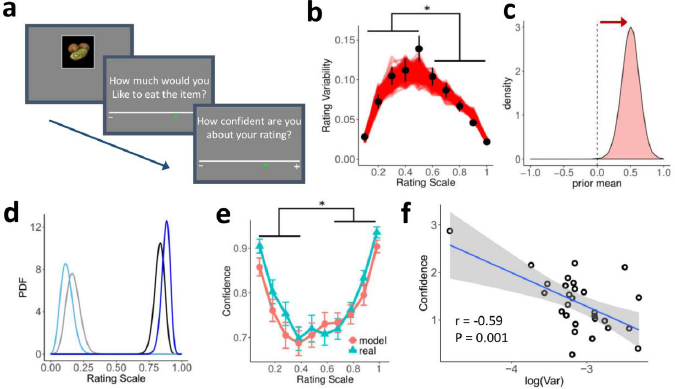
Confidence (Experiment 4). **a)** Participants provided value ratings as in experiments 1 and 2, but also rated their confidence in the value ratings. **b)** Rating variability plotted as a function of each item’s mean rating across both rounds (black dots; error bars represent s.e.m. across participants). We fitted the subjective-value-rating likelihood function (Eq. 1) to the empirical ratings. To test the quality of the model fits, we simulated 500 experiments. The simulated rating variability (semi-transparent red lines) was again higher for low-rated items (β= −0.18±0.04, P<0.001). **c)** Posterior estimates of the expected value of the prior are shifted towards higher rating values (the zero position maps to the center of the rating scale). **d)** Posterior densities were constructed for symmetric subjective values *v* (blue and black) in the unbounded subjective scale. Group of densities on the left (light colors) and right side (dark colors) reflect low and high subjective values, respectively. Given the expected value of the prior (panel c), the efficient coding model predicts lower levels of confidence for low-rated goods (relatively wide posteriors; light colors) than for high-rated goods (relatively narrower posteriors; dark colors). **e)** Empirical confidence ratings as a function of value ratings (blue) and predictions of the best-fitting confidence model (red; see also Supplementary Table 2). As predicted, confidence was higher for higher model-predicted subjective values *v_m_*(βRandEffects=0.22±0.070, P=0.002). **f)** Relationship of the partial effects between confidence and value rating variability (after controlling for rating value). As predicted by the model, these two metrics are negatively correlated (R = −0.59, probust=0.0011).

The data (see Figure 4e,f) confirm all three predictions. First, confidence increases for values closer to the extremes of the rating scale (quadratic effect β_RandEffects_=0.41±0.03, P<0.001). This replicates previous reports^15^ but now reveals the mechanistic origins of this effect in the efficient coding and Bayesian decoding framework. Second, confidence was higher for higher model-predicted subjective values (β_RandEffects_=0.22±0.070, P=0.002). This runs counter to previous suggestions^15^ that confidence ratings should be symmetric with respect to the center of the rating scale: Both our model and empirical data reveal that this is not necessarily the case, as the confidence ratings depend on the prior distribution (Supplementary Table 2). Third, across participants, the higher the variability in the subjective value estimations, the less confident the participants are (β_robust_ = −0.81±0.20, P<0.001). Crucially, this regression analysis controls for the mean value ratings for each participant (Fig. 4f), thereby confirming our model predictions that confidence relates to rating variability independently of how valuable the participants rated the items. Finally, to test these model predictions more quantitatively for the observed data using our framework, we implemented 11 different proposed models of how confidence may be derived^15,36–38^ from the posterior distributions of rating values given by our framework (see full details of the model specifications in Supplementary Table 2). We emphasize that for all these tests, we fixed both the parameters of the prior distribution *θ* and the subjective values *v(1,…,M)* for each participant to the values obtained by fitting the model to her prior rating data. We found that a model based on the statistical definition of confidence (i.e., the probability that the rating is correct^36,37^) provided the best fit to the empirical confidence reports (Fig 4e, Supplementary Table 2). This confirms that our efficient-coding model accurately captures the value inference processes that underlie not only subjective value estimates but also the reported confidence in these estimates.

## Discussion

Our work demonstrates that variability, biases, and confidence in preference-based choices are all consistent with information-maximizing transmission and statistically optimal decoding of values by a limited-capacity system. This suggests that principles governing the encoding and interpretation of low-level sensory signals are also relevant when humans report and choose based on subjective preferences. More specifically, our results support theoretical proposals according to which, just as in the case of sensory systems, subjective value systems optimize the use of limited resources for processing value information and exploit environmental regularities in order to optimally guide preference-based decisions^17,18,20^. Thus, our findings provide a fundamental step toward a unified account of how humans perceive and valuate the environment in order to optimally guide behavior.

Our work introduces a new framework that may serve to improve the modelling and prediction of preference-based decision making, and more generally any cognitive process that involves fully subjective value estimations (such as pain^39^ and health^40^ perception, to mention only two examples). We demonstrate that the common practice of using value ratings as inputs to decision models^11,13–15,29–31^ is suboptimal, since these reports should be just as variable and biased as the choices themselves that the experimenter wants to model. This is not a trivial issue, as both ratings and choices can be subject to the same complex non-linearities (Fig. 3) due to the encoding/decoding strategies implemented by the valuation system. Our model provides a solution to this problem, since it makes it possible to determine both the observer’s subjective values and their underlying prior distribution while accounting for capacity limitations with a single set of ratings. These parameters and values can then be used to predict fully independent preference-based decisions, with higher accuracy than existing standard approaches in the literature. This procedure obviously differs from traditional economic approaches that derive preferences directly from observed choices^41^ and ignore the processes involved in estimating subjective values (and the associated sources of variability). Our results show that this ignorance is not warranted; a detailed understanding of these processes should be a critical aspect to consider in theories of decision making and economic behavior^42^.

In our model specification, variability in subjective value estimates and choices emerges from both internal noise in the coding of value and unspecific (external) noise in the decision process. This perspective is fundamentally different from standard approaches where preference variability is solely attributed to unspecified noise in the decision process (e.g., noise added to deterministic value functions^16,43^ through the application of a softmax function). Based on our empirical and modelling work, we argue that positing unspecified sources of noise in the decision process may be insufficient, given that accounting for noise involved in the coding of value appears to be crucial for deriving more accurate predictions of economic decisions^22^. Even though some characteristics of the models used for this purpose (e.g., the exact loss function of the Bayesian encoder) may be refined by future research on both perception^44,45^ and subjective valuation, our findings clearly illustrate the general utility of this approach.

In some respects, our model of the coding of value resembles the one posited by decision-by-sampling theory^47^. That theory proposes that estimated subjective values directly reflect samples drawn from an internal noisy representation of value, but unlike in our analysis, no optimal Bayesian decoding is assumed. As a consequence, decision-by-sampling theory cannot account for our finding that biases in valuation are changed by time pressure, since drawing fewer samples should lead to value estimates that are noisier, but not different on average. This contrasts with the predictions of the framework presented here, which we show to be fully congruent with the empirically observed biases under time pressure.

Our results support the idea that reported confidence in subjective value estimations is well captured by a statistical measure of confidence^36,37^. While a recently proposed framework^15^ provides an elegant normative account of confidence judgments for individual value estimates, it does not provide a precise account of what information should actually be encoded but focuses only on what may be decoded. Additionally, that framework does not take into account the natural distribution of object values in the environment and capacity limitations in information processing. Our work provides a more comprehensive characterization, by demonstrating that the same efficient coding framework that accounts for biases and variability in subjective value estimates and choices also accounts for the reported confidence in these value estimates. In general, we hope that these results may motivate researchers to further develop explicit process models of metacognition^46^.

While our work highlights similarities between perceptual and value inference, it has remained agnostic as to how the internal response used for this purpose is derived from low- and high-level sensory signals. Understanding such feature extraction will be important for characterizing how the internal value response may be constructed, e.g. by sampling from memory^32,48^ and emotion systems^49^. While we formally demonstrate that the precision of encoded subjective values in capacity-limited systems may relate to the number of discrete samples that can be drawn (e.g., from memory or emotion systems), we have so far only focused on the encoding and readout of simple one-dimensional subjective values associated with an object. However, the framework used here could be parsimoniously extended to incorporate a whole range of lower-level sensory signals and to encompass more complex hierarchical structures. Despite this interesting challenge to further understanding the construction of preferences^10^, it is remarkable that a simple normative specification inspired by fundamental principles of low-level sensory perception can capture important aspects of preference-based decisions.

Bayesian models have often been criticized for allowing an arbitrary choice of prior and likelihood functions, as a consequence of which it is suggested that their predictions are vacuous^50^. However, in this study we have shown that by fully constraining the decision model to the distribution of object values – while taking account of capacity constraints – it is possible to accurately capture preference-based choice behavior using a parsimonious model. In line with previous work on low-level sensory perception^4,9^, our results demonstrate that the above-mentioned critique is not always valid. This should motivate researchers to pursue the identification of optimal solutions to computational problems posed by the environment – in both perception and subjective valuation – without ignoring the fact that biological systems are by definition limited in their capacity to process information.

Taken together, our findings suggest that resource-constrained models inspired by models of perception^3,9,17^ may have far-reaching implications not only in neuroscience,but also in psychology and economics^20,22,51,52^. Such models offer the prospect of explanations for seemingly irrational aspects of choice behavior, grounded in the need to represent the world with only finite precision. Recent work suggests that features of economic decisions such as risk aversion^22^ and preference reversals^20^ can be understood as further examples of biases resulting from optimal Bayesian inference from imprecise internal representations of value. This supports our emphasis on the desirability of developing models of decision making that account simultaneously for the goals of the organism, its environment, and its biological constraints.

## Methods

### Participants

The study tested healthy young volunteers (total n=127, age 19-37 years: n=38 in experiment 1, n=37 new participants in experiment 2 (replication of results obtained in experiment 1), n=24 new participants in experiment 3 and n=28 new participants in experiment 4). Participants were randomly assigned to each experiment. Sample size was determined based on previous studies using similar stimuli and tasks^11,53^. Participants were instructed about all aspects of the experiment and gave written informed consent. None of the participants suffered from any neurological or psychological disorder or took medication that interfered with participation in our study. Participants received monetary compensation for their participation in the experiment, in addition to receiving one food item in the decision-making task (see below). The experiments conformed to the Declaration of Helsinki and the experimental protocol was approved by the Ethics Committee of the Canton of Zurich.

For all experiments, participants were asked not to eat or drink anything for 3 hours before the start of the experiment. After the experiment, participants were required to stay in the room with the experimenter while eating the food item that they had chosen in a randomly selected trial of the decision-making task (see below). All experiments took place between 9am and 5pm.

### Value rating task

Experiments 1 and 2 consisted of three main phases: (1) rating phase 1, (2) rating phase 2, and (3) the decision-making task. In rating phase 1, we asked the participants to provide subjective preference ratings for a set of 64 food items using an on-screen slider scale (Figure 2a). All of the food items were in stock in our lab and participants were notified about this. Importantly, participants saw all food products before the ratings so that they could effectively use the full range of the rating scale. Moreover, participants knew that all products were randomly drawn from the two biggest supermarkets in Switzerland. Based on pilot measurements and previous studies^11,53^ in our lab, we selected food items that varied all the way from items that most participants would find unappealing (e.g., raw broccoli) to items that most participants would find highly appetitive (e.g., ice cream). This was important as our model should capture the full range of subjective values that humans typically assign to food items on a daily basis. During the ratings, participants indicated “how much they wanted to eat the presented food item at the end of the experiment”. The slider scale was continuous in experiment 1 with no numbers displayed (Fig. 2a), whereas the rating scale in experiment 2 was divided in 20 steps of equal size with numbers displayed under each step (Fig. 2a). This was done to ensure that the effects observed in Experiment 1 did not reflect the absence of reference points in the middle of the rating scale. Participants were informed that the rightmost endpoint would indicate items that they would most love to eat, whereas the leftmost endpoint would indicate items that they would most hate to eat. The initial location of the slider was randomized for each item to reduce anchoring effects.

Rating phase 2 was identical to rating phase 1 and took place immediately after phase 1. The order of the items’ presentation was randomized. Crucially, participants were not informed before the rating phase 1 that a second rating phase and a decision-making task would take place. This was important as it prevented participants from actively memorizing the location of the rating in the slider in the first phase, thus providing us with a clean measure of preference variability.

In Experiment 3, participants provided value ratings as in Experiment 1, but for the first round of ratings, half of the goods were presented with a duration of 900 ms and the second half with a duration of 2500 ms. For the second round of ratings, the presentation durations were inverted for each good. The exposure time (900 ms or 2500 ms) was pseudo-randomly selected for each good in the first round of ratings. Crucially, participants were not informed in advance of the time manipulations.

In Experiment 4, participants provided value ratings as in experiment 1, but indicated after each rating their confidence in their first-order rating (Fig. 4a). Following procedures of previous work^15^,we informed participants that the leftmost side of the rating scale means “Not at all” confident and the rightmost side means “Totally” confident.

### Choice task

For Experiments 1 and 2, immediately after the two rating phases, an algorithm selected a balanced set of decision trials divided into four value difference levels on the rating scale (rating difference ˜5%, ˜10%, ˜15% and ˜20% of the length of the rating scale), as defined by the average rating across phases 1 and 2 provided by each participant. Decision-making trials started with central presentation of a fixation cross for 1-2 seconds. Immediately after this, two food items were simultaneously displayed, one in the upper and one in the lower hemifield (Fig. 2a). The food items were presented until response and participants had up to four seconds to make a choice. Participants were instructed to choose which of the two items (upper or lower) they preferred to consume at the end of the experiment. To make these choices, participants pressed one of two buttons on a standard keyboard with their right-index finger (upper item) or their right thumb (lower item). In Experiments 1 and 2, we defined a consistent choice as a trial in which the subject chose the item with a higher mean rating from the prior rating phase. Each experimental session comprised a maximum of 240 trials (this depended on the rating distribution of each participant) divided into 6 runs of 40 trials each. The trials were fully balanced across rating-difference levels (˜5%, ˜10%, ˜15% and ˜20% of the length of the rating scale) and location of consistent response option (Up or Down).

## Model

We assume that the presentation of an object with stimulus value *v* elicits an internal noisy response *r* (*encoding*) that the observer uses to generate a subjective value estimate 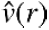 - the *decoded* stimulus value (Fig. 1a). For efficient value coding, a function *F*(*v*) maps the stimulus space to a new space where the Fisher information is uniform over the entire real line. This requires the definition

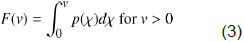

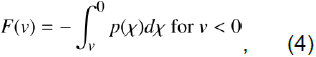

where *p(χ)* is an improper prior distribution, so that *F(v)→∞* as *v→∞*, and *F(v)→∞* as *v→-∞*. We assume that conditional on the value *v*, an internal noisy (neural) response is generated

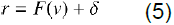

with δ˜*N(0,σ^2^)*, where σ measures the degree of noise in the internal representation that is constant over all possible values *F(v)*. We also note that the prior distribution for possible values *F(v)* of is uniform on the real line. The posterior mean estimate of *v* (the estimator that minimizes mean squared error) is then given by

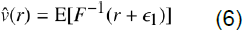

with ∊_1_˜*N(0,σ^2^)*. Here E[.] means the expectation over possible values of ∊_1_. This estimator is a deterministic function that maps each measurement *r* to an estimated subjective value 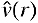; this deterministic mapping therefore cannot account for trial-to-trial fluctuation in the value estimates. The variability in the value estimates arises because of the variability in the measurement *r* on each trial. Accounting for this variability, it follows that for any true stimulus *v_0_*, the mean estimate should be given by

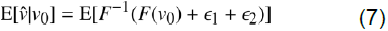

with ∊_2_˜*N(0,σ^2^)*. E[.] now means the expectation over possible values of (∊_1_+∊_2_)˜*N(0,2σ^2^)*. In the small-noise limit, we can take a second-order Taylor expansion:

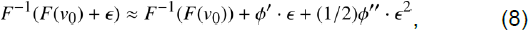

where 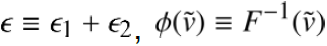, and the derivatives of 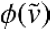 are evaluated at 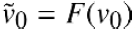. Taking the expected value over possible realizations of ∊_1_ we obtain

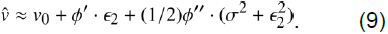

Conditional on a particular stimulus value *v_0_*, this is a random variable with expected value

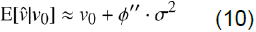

and variance

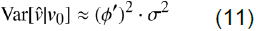

If we approximate this distribution by a normal distribution with the mean and variance just calculated above, we would obtain a probability density for given 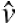 by

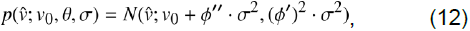

where *θ* is a set of parameters of the prior distribution (see below). This expression is the likelihood of a given subjective value estimate 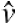conditioned on a true stimulus value. If one wants to write the joint likelihood of a pair of values 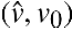 occurring when *v_0_* is drawn from the prior, one obtains

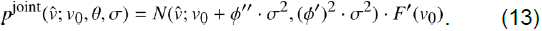

In addition to internal noise in the coding of value σ, we also account for late noise in the decision stage (i.e. post-decoding noise), which may capture any unspecific forms of downstream noise occurring during the response process that are unrelated to valuation per se. We assume this late noise to be normally distributed; it can therefore be easily added to our model as follows (see Supplementary Note 1 for further discussion on the different sources of noise)

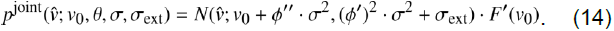

The last part of the model defines the probability distribution in the space of the bounded rating scale. Without loss of generality, we assume that this scale is bounded from 0 to 1 (this can obviously be adapted), with a monotonic mapping of subjective preference values that preserves preference ordering. Transforming the unbounded internal scale to this bounded physical scale requires a mapping that preserves monotonicity. A convenient and relatively simple function allowing this transformation is the logistic function: 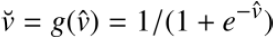, which provides a one-toone mapping of the estimate 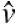 from the subjective to the physical scale in any given trial. The implied joint probability density 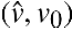 on the rating scale is thus given by

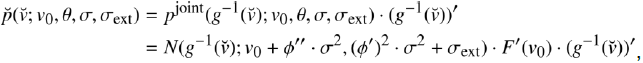

which is Equation 1 given in the main text. Here the inverse mapping of the subjective to the unbounded scale is given by

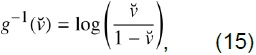

and its first derivative is

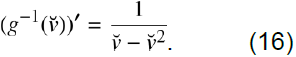

Recall that we are assuming here that the decision maker maximizes mutual information between the input stimulus and the noisy measurement, therefore *F(v)* is defined as the cumulative density function (CDF) of the prior distribution *p(v)*. Here we assume that the prior follows a logistic distribution

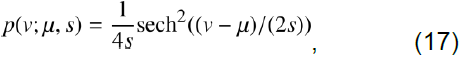

where *μ* and *_s_* represent the mean and scale, respectively. The advantage of using this distribution is that its CDF and both the first and the second derivative of the quantile function have closed form solutions; however, any similar prior distribution could be used without greatly affecting the quantitative predictions presented here.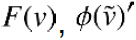, and 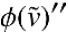are given by:

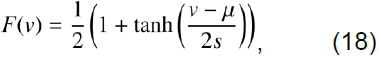

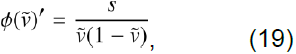

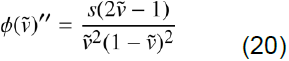

Thus, for any experimental data set consisting of *M* goods and *N* value ratings for each good, we can find the set of parameters of the prior, the internal valuation noise σ, external noise σ_ext_, and the “true” stimulus values *v(1,…,m)* that maximize the likelihood of the observed set of ratings (under the constraint that *v(1,…,m)* is distributed following *P(v)*).

In order to compute choice consistency predictions that an experimenter would obtain when performing such analysis in the rating scale (Fig. 1b,c), we first computed for a fine-grained sequence of subjective values *v_0_* their corresponding expected value and variance perceived in the rating scale (if we assume that the experimenter can obtain a large number of ratings for each good) via

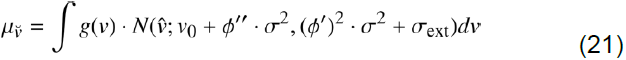

and

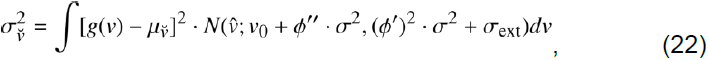

where the subscript (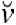) reflects the expected value (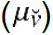) and variance (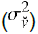) at position 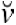in the rating scale. We then looked for expected values 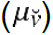closer to the values [0.1,0.11,0.12,…0.89,0.9] and used their corresponding variance to approximate the level of choice consistency as follows:

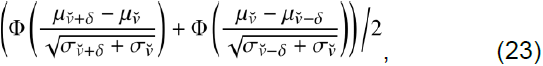

with *δ* = 0.05 (note that different values of *δ* move the accuracy curve up or down but does not affect the general u-shaped curve obtained in our predictions; see Fig. 1b,c).

## Behavioral analyses

Preference-rating variability in Experiments 1,2 and 4 was computed as the standard deviation (SD) for each item across the rating phases 1 and 2. To visualize this effect, we plotted the SD as a function of the mean rating (Figs. 2b,c; 4b). To investigate the influence of both VD and rating variability on the consistency of choices (Fig. 2d), we performed a hierarchical logistic mixed-effects regression of choices (defining consistent=1, inconsistent=0) on our three main regressors of interest, namely: value difference (VD), summed-variability (Var, defined as the sum of the two SDs of the two food items presented in each trial), and the summed-value (OV, defined as the sum of mean rating values of the two food items presented in each trial). All regressors of interest were included in the same model. Similarly, all the population-level regressions described for Experiment 4 were based on a hierarchical linear mixed-effects regression approach. All mixed-effects regressions in this study had varying subject-specific constants and slopes (i.e., random effects analysis). Posterior inference of the parameters in the hierarchical models was performed via the Gibbs sampler using the Markov Chain Monte Carlo (MCMC) technique implemented in JAGS^54^, assuming flat priors for both the mean and the noise of the estimates. For each model a total of 10,000 samples were drawn from an initial burn-in step and subsequently a total of new 10,000 samples were drawn with three chains (each chain was derived based on a different random number generator engine, and each with a different seed). We applied a thinning of 10 to this final sample, thus resulting in a final set of 1,000 samples for each parameter. We conducted Gelman–Rubin tests^5^5 for each parameter to confirm convergence of the chains. All latent variables in our Bayesian models had 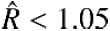, which suggests that all three chains converged to a target posterior distribution. The “p-values” reported for these regressions are not frequentist p-values but instead directly quantify the probability of the reported effect differing from zero. They were computed using the posterior population distributions estimated for each parameter and represent the portion of the cumulative density functions that lies above/below 0 (depending on the direction of the effect). The regressions across participants reported for Experiments 1,2 and 4 were computed using robust linear regressions using the rlm function^56^ implemented in the statistical computing software R^57^.

In order to fit the efficient coding model to the rating data in Experiments 1, 2 and 4, we found the stimulus values *v(1,…,M)*, parameters of the prior *θ*, encoding noise σ and external noise σ_ext_ that maximized the likelihood function 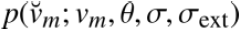 (Eq. 1 in main text) of the observed set of ratings for each participant under the constraint that *v(1,…,M)* is distributed following *P(v;θ)*(Supplementary Fig. 3; methods). Alternatively, defining 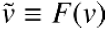, one can find the values estimates in the efficient space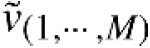 under the constraint that these are uniformly distributed. Using either approach, we found nearly identical results for the fitted parameters, which is expected for correct model specification. Posterior inference of the parameters for this model can be conveniently performed via the Gibbs sampler.

We used the stimulus values *v(1,…,M)* and prior parameters *θ* fitted to the rating in order to predict choices in the two-alternative choice task in Experiments 1 and 2. Following our modelling specification, over many trials the probability that an agent chooses an alternative with stimulus value *v_1_* over a second alternative with stimulus value *v_2_* is given by:

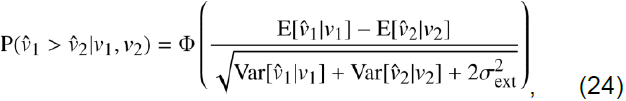

where Ф() is the CDF of the standard normal distribution the expressions for 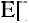 and 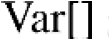 are given in Eqs. 10 and 11 (see above). In other words, the input values of the choice model are fully constrained by the efficient coding model and therefore the choice model has only n=2 free parameters, namely the resource noise of the encoder σ and the external noise σ_ext_. Fits to the choice data were performed via the Gibbs sampler using a hierarchical Bayesian model assuming flat priors for both noise terms. When evaluating different models, we are interested in our model’s predictive accuracy for unobserved data, thus it is important to choose a metric for model comparison that considers this predictive aspect. Therefore, in order to perform model comparison, we used a method for approximating leave-one-out cross-validation (LOO) that uses samples from the full posterior^58^. The smaller the LOO the better the fit. We found that in Experiments 1 and 2, the best model was the efficient-coding model. Crucially, this finding is fully replicated when using a different Bayesian metric such as the wAIC^58^. Description of the different choice models tested here is presented in Supplementary Figure 6.

## Acknowledgements

R.P. thanks Xue-Xin Wei and Alan Stocker for inspiring discussions. This work was supported by a grant of the Swiss National Science Foundation (grant IZK0Z1_173607) and an ERC starting grant (ENTRAINER) to R.P; by grants of the Swiss National Science Foundation (grants 105314_152891 and 100019L_173248) and an ERC consolidator grant (BRAINCODES) to C.C.R; and a grant of the U.S. National Science Foundation to M.W.

## Data and code availability

The data that support the findings of this study and the analysis code are available from the corresponding author upon request.

## Author contributions

R.P and C.C.R. designed the study. R.P. collected and analyzed the data. All authors interpreted the results and wrote the manuscript.

